# GRETA: an R package for mapping *in silico* genetic interaction and essentiality networks

**DOI:** 10.1101/2022.09.21.508787

**Authors:** Yuka Takemon, Marco A. Marra

## Abstract

**Summary:** Mapping genetic interaction and essentiality networks in human cell lines have been used to identify vulnerabilities of cells carrying specific genetic alterations and to associate novel functions to genes, respectively. *In vitro* and *in vivo* genetic screens to decipher these networks are resource-intensive, limiting the throughput of samples that can be analyzed. In this application note, we provide an R package we call Genetic inteRaction and EssentialiTy mApper (GRETA). GRETA is an accessible tool for *in silico* genetic interaction screens and essentiality network analyses using publicly available data, requiring only basic R programming knowledge.

**Availability and implementation:** The R package, GRETA, is licensed under GNU General Public License v3.0 and freely available at https://github.com/ytakemon/GRETA and https://doi.org/10.5281/zenodo.6940757, with documentation and tutorial. A Singularity container is also available at https://cloud.sylabs.io/library/ytakemon/greta/greta.

**Supplemental information:** Supplemental materials are available at *Bioinformatics* online.

**Issue section:** *Systems biology*

## Introduction

Genetic perturbation screens, such as CRISPR-Cas9 knockout (KO) screens, have been used to uncover lethal and alleviating genetic interactions (GIs; Wang *et al*., 2017; Shalem *et al*., 2014; Hart *et al*., 2015; Behan *et al*., 2019) and essentiality networks to decipher a gene’s functional membership (Pan *et al*., 2018; Wainberg *et al*., 2021). Two genes are said to share a lethal, or synthetic lethal (SL), GI when the perturbation of both genes together in the same cell impedes cell viability, but perturbation of either gene individually has no impact on viability (Mani *et al*., 2008). Conversely, an alleviating GI occurs when perturbation of two genes leads to a fitness advantage over cells with either gene perturbed individually (Mani *et al*., 2008). Lethal GIs of genes mutated in cancers have revealed tumor-specific vulnerabilities that can be therapeutically exploited. Most notable are the lethal GIs between *BRCA1/2* and *PARP1* (Bryant *et al*., 2005; Farmer *et al*., 2005), which led to the use of PARP inhibitors in BRCA1/2-deficient breast (Robson *et al*., 2017, 2019), ovarian (Mirza *et al*., 2018), and prostate cancers (Ramakrishnan Geethakumari *et al*., 2017). Essentiality network maps have been utilized to assign novel functions to genes (Wang *et al*., 2017; Kim *et al*., 2019; Wainberg *et al*., 2021; Pan *et al*., 2018). Two genes share co-essentiality when perturbation of either gene leads to similar fitness effects in cells. This is interpreted to indicate that the genes may have similar or synergistic functions (Kim *et al*., 2019; Wainberg *et al*., 2021). In contrast, two genes are anti-essential when perturbations in either of them lead to opposite cellular fitness effects. This is interpreted to indicate genes with inhibitory or antagonistic relationships (Kim *et al*., 2019; Wainberg *et al*., 2021). It seems clear that mapping GI and essentiality networks of cancer-associated genes could reveal biological functions that might be leveraged to inform potential therapeutic strategies.

The Cancer Dependency Map (DepMap) public data platform (Ghandi *et al*., 2019; Meyers *et al*., 2017) the largest resource for mining GI and essentiality networks of human cancer cell lines. DepMap contains data from multi-omic characterization of cancer cell lines, including whole-genome sequencing, exome sequencing, RNA sequencing (RNA-seq) (Ghandi *et al*., 2019), proteomic quantification (Nusinow *et al*., 2020), and genome-wide CRISPR-cas9 KO screens (Meyers *et al*., 2017), which are updated bi-annually with additional cancer cell lines (detailed in https://depmap.org/portal/announcements/). Novel algorithms utilizing DepMap data to predict GIs have emerged (De Kegel *et al*., 2021; Chiu *et al*., 2021; Benfatto *et al*., 2021). However, these tools either lack flexibility, the ability to detect alleviging GIs, or require advanced programming knowledge. Thus, for various reasons, these tools have limited accessibility to the larger research community.

To facilitate discoveries using GI and essentiality network mapping, we created an R package we called Genetic inteRaction and EssentialiTy mApper (GRETA), which leverages the DepMap data platform. GRETA provides a simplified approach to query DepMap data, and provides user-driven selection of cell line features, including by gene, mutation, and cancer types, to predict both lethal and alleviating GIs and essentiality networks. GRETA provides intuitive visualization of results that requires only a basic understanding of the R programming language.

## Materials and Methods

### *Performing* In Silico *GI Screens*

GRETA leverages cell line annotations, mutation annotations, copy number quantifications, RNA-seq, proteomic quantifications, and genome-wide CRISPR-Cas9 KO screen data that were made available by the DepMap data platform (Supplemental Methods; Meyers *et al*., 2017; Ghandi *et al*., 2019; Nusinow *et al*., 2020) and predicts candidate GIs in three steps: 1) selecting control and mutant cancer cell lines; 2) surveying mRNA and/or protein expression changes; and 3) predicting and visualizing GIs (illustrated in Supplemental Figure S1A). Briefly, the first step is driven by a user-defined feature, such as a gene of interest (GOI). By default, when a GOI is provided, all pan-cancer cell lines with wildtype (WT) alleles and neutral copy number of the GOI are assigned to the control group, and lines with loss-of-function (LOF) alterations in the GOI are assigned to one of the mutant subgroups, including homozygous deletion (HomDel), *trans*-heterozygous deletion (T-HetDel), and heterozygous deletion (HetDel) groups (see Supplemental Methods for details). Cell lines can also be selected by specific mutation, cancer type, or manual curation (Supplemental Methods).

The second step involves extracting DepMap mRNA and protein expression data (if available) and performing differential mRNA and protein expression analyses of the GOI between control cell lines and mutant cell lines (Supplemental Methods). A mutant group with a significant reduction in mRNA or protein expression compared to the control group by Welch’s T-test is consistent with potential reduction of GOI functions, and thus a user may select such a mutant cell line group for downstream analysis.

Finally, candidate GIs are predicted by conducting differential lethality probability analyses, which identify a gene KO that results in opposing survival outcomes in the mutant cell lines and control cell lines. Mann-Whitney U tests are performed by extracting all lethality probabilities from the DepMap CRISPR-cas9 KO screen data and comparing the control and mutant cancer cell line groups. GRETA also generates an interaction score that is used to visualize candidate GIs and to rank them from most to least likely to be lethal or alleviating (Supplemental Methods).

### Mapping Essentiality Networks

GRETA adapted a previously published Pearson correlation-based method for mapping essentiality between genes (Wang *et al*., 2017; Pan *et al*., 2018; Kim *et al*., 2019; illustrated in Supplemental Figure S1B; Supplemental Methods). Using this method, co-essential (positively correlated) and anti-essential (negatively correlated) genes can be identified. Ranked correlation coefficients are then used to visualize essentiality networks.

## Case study: Predicting pan-cancer GIs and co-essentiality networks of *ARID1A*

*ARID1A* is a member of the mammalian switch/sucrose non-fermentable (SWI/SNF) complex (Mashtalir *et al*., 2018) with a known SL GI to its paralog *ARID1B* (Helming *et al*., 2014). Rationalizing that this known SL interaction and complex association would serve as positive controls, we applied GRETA to predict pan-cancer GIs and essentiality networks of ARID1A, expecting to replicate known interactors and co-essential genes, respectively. We identified an *ARID1A* WT control group and an *ARID1A* HomDel mutant group, consisting of 529 and 13 cell lines, respectively. The *ARID1A* HomDel mutant group showed significantly reduced *ARID1A* mRNA expression compared to the control group (Welch’s t-test p-value < 0.001; Supplemental Figure S2A-B; Supplemental Table S1A-B), consistent with potentially reduced ARID1A function. Using these two groups, we found that *ARID1B* was predicted to be the top lethal genetic interactor of *ARID1A* (Supplemental Figure S2C; Supplemental Table S1C), thus confirming the established SL GI between these two genes. Furthermore, a pan-cancer essentiality analysis of *ARID1A* revealed the expected top five list of co-essential genes that are known mSWI/SNF complex members, namely *ARID1A, SMARCE1, SMARCB1, SMARCC1*, and *DPF2* (Mashtalir *et al*., 2018; *Supplemental Figure S2D; Supplemental Table S1D). GRETA was thus able to recapitulate known GIs and co-essential networks of ARID1A*.

## Discussion

GRETA was built to provide an easy-to-use method for mapping GI and essentiality networks. The R package requires only a basic understanding of the R programming language and we provide an easy-to-follow tutorial (https://github.com/ytakemon/GRETA). Using GRETA, we replicated known GIs and essentiality networks of *ARID1A*, thus demonstrating how the tool can be used and ultimately applied to discover GI and essentiality networks of other GOI. We are currently aware of 115 visitors and 52 total unique installations of GRETA and from this deduce that GRETA is of interest to the wider research community.

## Data Availability Statement

The GRETA R package, data analyzed in this study (DepMap version 20Q1), and the code used to generate the supplemental figures and tables are publicly available as a tutorial provided on the GRETA GitHub repository (https://github.com/ytakemon/GRETA). GRETA has been archived with a citable DOI on Zenodo (https://doi.org/10.5281/zenodo.6940757) and a Singularity container has been made available on Sylabs (https://cloud.sylabs.io/library/ytakemon/greta/greta) to ensure reproducibility. GRETA is shared under the GNU General Public License v3.0.

## Acknowledgments

We would like to acknowledge the Broad Institute for generating and making their DepMap data platform available for the research community. We would like to thank Drs. Anindya Dutta and Yoshiyuki Shibata at the University of Alabama at Birmingham Marnix E. Heersink School of Medicine for testing GRETA and for the discussions that helped improve its documentation. We would also like to thank Richard Corbett at Canada’s Michael Smith Genome Sciences Centre for the discussion and guidance on containerizing the tool for reproducibility.

