## Supplemental Methods for "GRETA: an R package for mapping *in silico* genetic interaction and essentiality networks"

*Human Cancer Cell Line -omic and Genetic Screening Data*

GRETA is compatible with DepMap data versions from v20Q1 to the latest version, which contains cell line annotations, mutation annotations, copy number quantifications, RNA-seq, proteomic quantifications, and genome-wide CRISPR-Cas9 KO screen data. All necessary DepMap data can be easily obtained as Rdata files from the GRETA GitHub repository and is compliant with Broad Institute's terms of use. Alternatively, data may be sourced from DepMap's figshare (https://figshare.com/authors/Broad_DepMap/5514062).

*Querying Cell Line by Mutation and Context*

Based on a user-defined GOI, a series of `list_available` functions are provided to query and summarize mutations that are present in all pan-cancer DepMap cancer cell lines. Optionally, these queries can be filtered further by primary cancer types and subtypes provided by DepMap. To perform this operation, GRETA leverages the following DepMap data files: `CCLE_mutations.csv`, a mutation annotation format (MAF) file, for mutation annotations; and `sample_info.csv` for cell line context information.

*Identifying Cancer Cell Lines of Interest*

Once a GOI and context are determined, control and GOI mutant cancer cell lines are selected using `select_cell_lines()`. By default, GRETA will select mutant cell lines with LOF alterations and copy number alterations affecting the GOI and control cell lines that carry the WT GOI allele. LOF alterations are queried from the DepMap MAF file and include variants annotated as nonsense mutation, frameshift insertion and deletion, splice site mutation, and start codon insertion and deletion. Zygosity of an alteration is also inferred from the allele count columns provided in the MAF file. An alteration with zero counts generated from the reference allele is considered a homozygous alteration, and all other allele counts are considered heterozygous alterations. Copy number alterations annotation utilizes DepMap's `CCLE_gene_cn.csv` file, which contains relative gene-based $log_2$-transformed copy number counts. Cell lines are annotated as either having neutral copy number (>= 0.75 & < 1.25 relative copy number counts), shallow loss (> 0.25 & < 0.75 relative copy number counts), deep deletion (<= 0.25 relative copy number counts), or amplified copy number (>= 1.25 relative copy number counts) of a GOI. In addition to the control cell line group described in the main text, cell lines are categorized into one of the following mutant subgroups HomDel, T-HetDel, HetDel, Amplified, and Other. HomDel cell lines harbor homozygous LOF alterations or have deep copy number loss; T-HetDel lines are possible *trans*-heterozygotes that harbor two or more heterozygous LOF alterations with neutral or one LOF mutation and shallow copy number loss; HetDel lines contain a single heterozygous LOF alteration with neutral copy number or no LOF alterations with shallow copy number loss; Amplified lines harbor no LOF alterations in the GOI but do carry amplified copy number; and the Other group contains lines that harbor both LOF alterations in and amplified copy number of the GOI.

*In Silico Genetic Screening*

GI prediction is performed using `GI_screen()`, which conducts a Mann-Whitney U-based differential lethality probability analysis to compare the lethality probability of all targeted genes (provided in the `CRISPR_gene_dependency` or `Achilles_gene_dependency.csv` DepMap file, depending on the data version) between the control and mutant cancer cell line groups. This generates a p-value, which is then used to compute the Benjamini-Hochberg

(BH)-adjusted p-value for each targeted gene (Benjamini and Hochberg, 1995). Furthermore, GRETA provides additional metrics such as the median, mean, standard deviation, and $\log_2$ fold changes of the median and mean lethality probabilities for each gene in control and mutant group cell lines. The tool also provide Cliff's delta for a non-parametric effect size between control and mutant lethality probabilities (Cliff, 1993), and a Hartigan's dip test p-value to indicate whether a multimodal lethality probability distribution was observed to identify whether confounding factors may be present (Hartigan and Hartigan, 1985). In order to rank lethal and alleviating GIs, GRETA generates an interaction score for each gene screened (represented as $i$); where $\rho_i$ represents the Mann-Whitney U test p-value and $\eta i_{mut}$ and $\eta i_{ctrl}$ represent the median lethality probabilities of the mutant group and control group, respectively:

$$interaction\ score = \ -\log_{10}\left(\rho_i\right) \times \sin\left(\log_2\left[\frac{\eta i_{mut}}{\eta i_{ctrl}}\right]\right)$$

A positive interaction score indicates lethal GIs and negative scores indicate alleviating GIs. By default, our cutoff for candidate GIs is set to a p-value < 0.05 and a median lethality probability of 0.3 in at least one cell line group. Finally, a ranked interaction score plot can be generated using `plot_screen()`, with the top user-defined number of lethal and alleviating GIs annotated on the plot (Supplemental Figure S1A).

*Essentiality Analysis*

GRETA utilizes DepMap's `CRISPR_gene_effect.csv` or `Achilles_gene_effect.csv` file (depending on the data version) containing KO effect scores for each gene targeted by the CRISPR-Cas9 screen to perform the essentiality analysis. `coessential_map()` uses a previously established method to discover co-essential and anti-essential genes of a GOI (Wang *et al.*, 2017;

Pan *et al.*, 2018; Kim *et al.*, 2019). This function calculates the Pearson correlation coefficient and p-value between the KO effect score of the GOI and all genes targeted in the screen. Furthermore, `get_inflection_points()` utilizes the RootsExtremalInflections R package (Christopoulos, 2019) to calculate the inflection points of the positive and negative coefficient curves. To objectively determine a threshold, we consider genes with a correlation coefficient above the positive inflection point or below the negative inflection point and with a p-value < 0.05 as candidate co- or anti-essential genes, respectively. The results can be visualized as a ranked essentiality plot using `plot_coessential_genes()`, with a user-defined number of top co- and anti-essential genes labeled (Supplemental Figure S1B).
