## Supplemental Figures and Table legend for "GRETA: an R package for mapping *in silico* genetic interaction and essentiality networks"

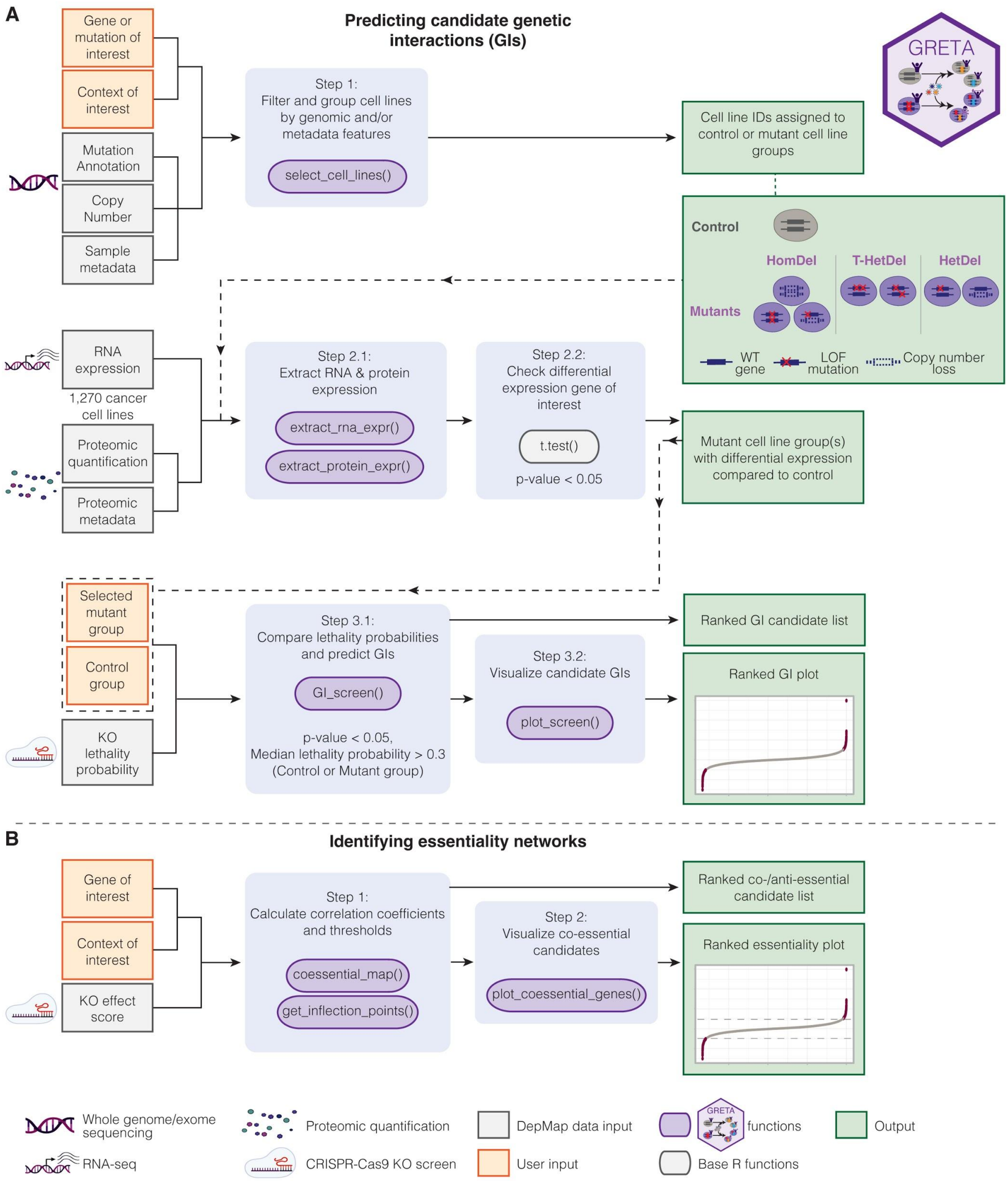


**Supplemental Figure S1.** A workflow of two GRETA package analysis modes. A) A four-step workflow to predict and visualize candidate GIs of a user-defined gene and/or cell feature of interest. Final outputs consist of a comma-separated (csv) file containing a ranked list of candidate genes that are potential GIs of the gene and/or cell feature of interest, and a ranked scatter plot with top lethal and alleviating GIs labeled for visualization. B) A two-step workflow to identify co-essential and anti-essential genes of a user-defined gene and/or cell feature of interest. Final outputs consist of a .csv file containing a ranked list of co-essential and anti-essential genes, and a ranked scatter plot to quickly visualize genes that are most likely to share essentiality. Icons from BioRender.com were incorporated in this figure.

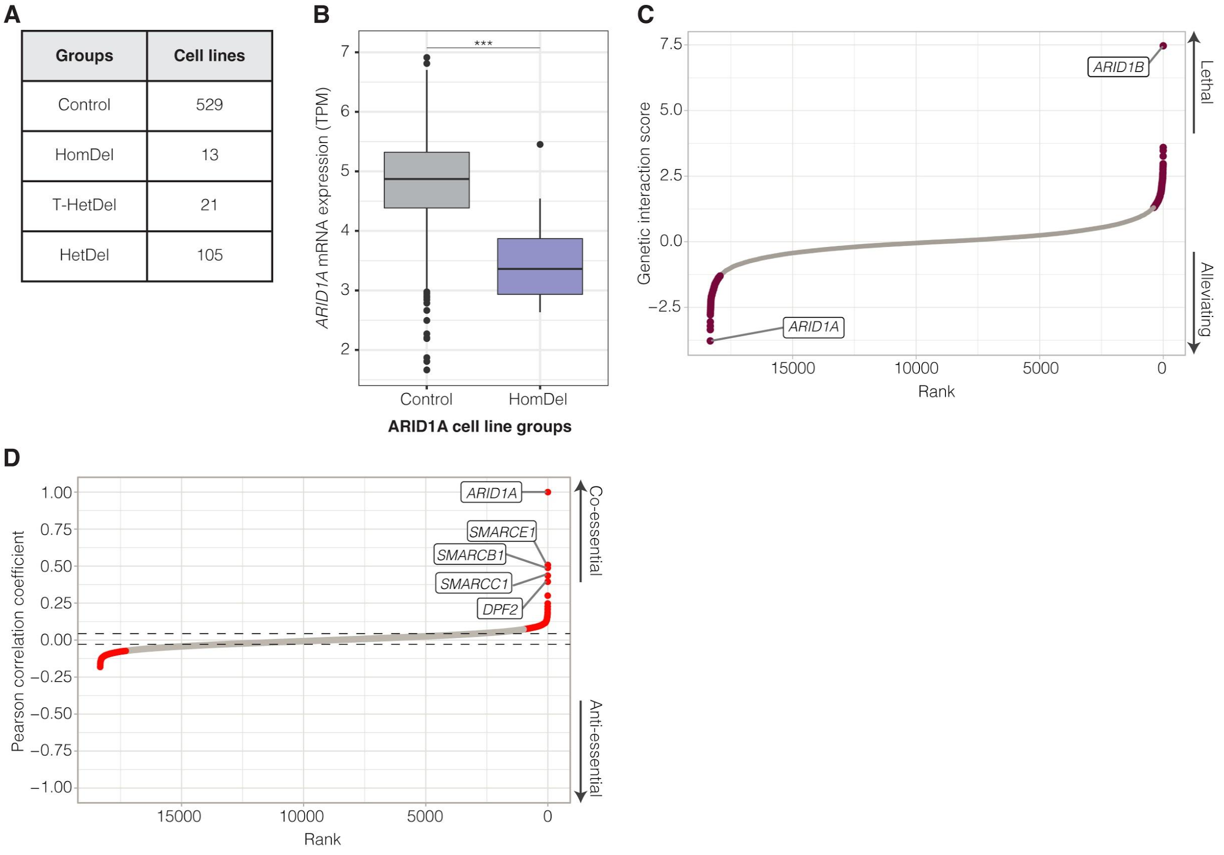


**Supplemental Figure S2.** Predicting *ARID1A* GIs and co-essential genes. A) Number of cancer cell lines in *ARID1A* control and LOF mutant groups identified using default pan-cancer settings. B) Tukey boxplots of *ARID1A* mRNA expression in *ARID1A* control (525 lines) and HomDel mutant groups (13 lines). Welch's T-test p-value *** < 0.001. C) Ranked *ARID1A* genetic interaction scores generated using GRETA. Purple points indicate candidate *ARID1A* GIs (152 total candidate genes, of which 102 were alleviating interactors and 50 were lethal interactors; Mann-Whitney U-test p-value < 0.05). The top most lethal and alleviating genetic interactors are labeled. D) A ranked *ARID1A* coefficient plot showing candidate co-essential genes and anti-essential genes (red points; Pearson correlation p-value < 0.05 and past inflection point of the positive or negative curve) generated using GRETA. Dashed horizontal lines denote the inflection points of the positive and negative correlation curve. The top five co-essential genes were labeled.

**Supplemental Table Legend**

**Supplemental Table S1.** GRETA outputs of *ARID1A in silico* GI screen and essentiality mapping case study. A) Pan-cancer *ARID1A* control and mutant cell line groups identified using default settings of `select_cell_lines()`. B) *ARID1A* mRNA expression retrieved for *ARID1A* control (525 lines) and HomDel mutant groups (13 lines) using `extract_mrna_expr()`. C) Output of `GI_screen()` comparing *ARID1A* control and HomDel cancer cell lines showing candidate lethal and alleviating GIs of *ARID1A*. D) Output of `coessential_map()` showing *ARID1A*'s pan-cancer candidate co-essential and anti-essential genes.
